# HortGenome Search Engine, a universal genomic search engine for horticultural crops

**DOI:** 10.1101/2024.01.01.573844

**Authors:** Sen Wang, Shangxiao Wei, Yuling Deng, Shaoyuan Wu, Haixu Peng, You Qing, Xuyang Zhai, Shijie Zhou, Jinrong Li, Hua Li, Yijian Feng, Yating Yi, Rui Li, Hui Zhang, Yiding Wang, Renlong Zhang, Lu Ning, YunCong Yao, Zhangjun Fei, Yi Zheng

## Abstract

Horticultural crops comprising fruit, vegetable, ornamental, beverage, medicinal and aromatic plants play essential roles in food security and human health, as well as landscaping. With the advances of sequencing technologies, genomes for hundreds of horticultural crops have been deciphered in recent years, providing a basis for understanding gene functions and regulatory networks and for the improvement of horticultural crops. However, these valuable genomic data are scattered in warehouses with various complex searching and displaying strategies, which increases learning and usage costs and makes comparative and functional genomic analyses across different horticultural crops very challenging. To this end, we have developed a lightweight universal search engine, HortGenome Search Engine (HSE; http://hort.moilab.net), which allows querying genes, functional annotations, protein domains, homologs, and other gene-related functional information of more than 400 horticultural crops. In addition, four commonly used tools, including ‘BLAST’, ‘Batch Query’, ‘Enrichment analysis’, and ‘Synteny Viewer’, have been developed for efficient mining and analysis of these genomic data.

## Introduction

Horticultural crops comprise fruits, vegetables, floricultural and ornamental plants, as well as beverage, medicinal and aromatic plants, and have played critical roles in food supply, human health, and beautifying landscapes. With the growing human population, new demands are placed on the yield, quality, diversity, and nutritional value of horticultural crops. Decoding the genomes of horticultural crops not only provides an opportunity to investigate gene functions and regulatory networks^1,2^, but also serves as the cornerstone for functional and comparative genomics studies^3,4^ and paves a path to resolve complex QTLs of important horticultural traits^5^. Advanced genome editing technologies have been demonstrated in recent years to have a great potential for improving the quality and yield of horticultural crops^6^, and reference genomes provide precise sequences for the application of genome editing technologies. Thus, genome sequencing plays a crucial role in horticultural crop improvement, and serves as an important foundation for understanding the history of crop domestication and evolution.

With the rapid advances of sequencing technologies, especially the PacBio HiFi long-read sequencing technology, various horticultural crop genomes have been deciphered, including those with high heterozygosity and polyploidy levels. To store, mine, and analyze the large-scale genomics data of horticultural crops, numerous databases have been developed, such as Sol Genomics Network (SGN), Genome Database for Rosaceae (GDR), Cucurbit Genomics Database (CuGenDB), among others^7–10^. Most of these databases manage genomic data for plants from a single-family or species^11^. Therefore, the genomic resources of horticultural crops are scattered in different databases, and these databases exhibit different ways of presenting and utilizing results, resulting in certain difficulties for users, especially in terms of searching tools that differ in complexity and functionality. This creates a learning curve for users seeking to search, browse, and conduct comparative analysis of genomic data across a broader range of plant species.

In recent years, there has been an increasing focus on using search engines to explore the genetic makeup of plants^12^. This has proven to be an invaluable tool for researchers who are interested in studying plant genomics, functional genomics, and molecular assisted breeding. To this end, we have developed the HortGenome Search Engine (HSE; http://hort.moilab.net), a lightweight universal search engine for the genomic data of horticultural crops. Compared to other genomic databases, it stands out for its search engine-like interface that allows users to easily search genomic data without requiring prior knowledge. Currently, the searchable genomic data includes species information, gene sequences, comprehensive functional annotations, and homologous gene pairs. The HortGenome Search Engine contains data of 434 genome assemblies for horticultural crops covering fruit trees, vegetables, ornaments, and beverage plants, as well as model plant species, Arabidopsis and rice. In addition to the searching function, several commonly used genomic data mining and analysis tools have been implemented in HSE, including ‘BLAST’, ‘Batch Query’, ‘Enrichment analysis’, and ‘Synteny Viewer’.

## DATABASE CONTENTS AND FEATURES

### Preparation of genomic data

More than 1000 genome assemblies of nearly 800 plant species have been sequenced and published by the end of 2021^13,14^. Genomic data of horticultural crops, including the genome sequences, gene structure annotations in general feature format (GFF), and mRNA, coding (CDS) and protein sequences of protein-coding genes, were collected from plant genomics, comparative genomics, and plant family-specific databases, such as Phytozyme^15^, Ensembl Plants^16^, Genome Warehouse in National Genomics Data Center^17^, SGN^7^, GDR^8^ and others. For some genome assemblies, only the genome sequences and GFF files are available; therefore, the corresponding mRNA, CDS and protein sequences were extracted using the gffread program^18^. We further performed quality control on the collected genomic data to ensure the accuracy of the data to be included in the database. For example, genome assemblies that lack a GFF file or have an inaccurate GFF file in which the numbers of genes or gene IDs were inconsistent with the corresponding mRNA, CDS and protein sequence files, were excluded. Finally, a total of 434 genome assemblies for horticultural crops, as well as the model plant species Arabidopsis and rice, were collected and included in the database (Table S1). Besides the genomic data, the taxonomy information, statistics of genome assemblies, associated publications, and images of the plant species, have also been collected from the PlaBiPD database (https://www.plabipd.de/), published manuscripts, and other data sources, and included in the database.

### Gene functional annotation

We used the pipeline described in our previous studies^9,19^ to generate comprehensive functional annotations for all protein-coding genes of the collected genome assemblies of horticulture plants. Briefly, protein sequences of the predicted genes were blasted against the GenBank non-redundant (nr), UniProt (TrEMBL and SwissProt), and Arabidopsis protein databases using DIAMOND^20^ with an E-value cutoff of 1e-4. Based on the identified homologs from the UniProt and Arabidopsis protein databases, concise and informative functional descriptions were assigned to each gene using the AHRD program (https://github.com/groupschoof/AHRD). Protein sequences were further compared against the InterPro database using InterProScan^21^ to identify functional protein domains. Transcription factors (TFs), transcriptional regulators (TRs), and protein kinases (PKs) were identified using the iTAK pipeline^22^.

To generate GO and KEGG pathway annotations for functional enrichment analyses, protein sequences were compared against the EggNOG database using eggnog-mapper^23^. The assigned GO terms of genes/transcripts retrieved from the eggnog-mapper results were converted to the GO Annotation File (GAF) format. In the eggnog-mapper results, some non-plant KEGG pathways were assigned to plant genes/transcripts. For example, the tomato gene *Solyc09g008400*, which encodes a serine/threonine protein phosphatase 2A regulatory subunit protein, was assigned to map05165, the human papillomavirus infection pathway. These non-plant pathways were manually identified and removed from the eggnog-mapper results.

### Synteny blocks and homologous gene pairs

Identifying synteny blocks and homologous gene pairs within or across genomes lays the groundwork for discovering and dating ancient genomic evolution events, as well as for inferring gene functions^24^. Detection of synteny blocks among all the 434 genomes would yield more than 90,000 pairwise genome comparisons, which is time-consuming and computationally not feasible. Therefore, in our study syntenic blocks and homologous gene pairs were identified only between any two genome assemblies from species within the same family, and within each genome assembly. In addition, synteny blocks and gene pairs were also identified between any genome assemblies and their corresponding model plants, i.e., Arabidopsis for eudicot plants and rice for monocot plants. Briefly, the CDS of each genome were arranged in the order based on the GFF file, and then the CDS from different chromosomes, linkage groups, or scaffolds of the two compared genomes were aligned using the LASTZ program with default parameters. Syntenic blocks and homologous gene pairs were then identified using the python version of MCScanX^25^, which implements a new BLAST filter to remove weak syntenic regions and tandem duplications^24^. In the end, a total of 1,832,351 syntenic blocks and 413 million homologous gene pairs were identified from 6,994 pairwise genome comparisons and imported into the back-end database.

### Data integration and indexing

Genome sequences, gene structures, and functional descriptions are imported into MongoDB, a popular NoSQL document database (https://www.mongodb.com/). Currently the database contains more than 34 million records of genes and transcripts from 434 genome assemblies. The top BLAST hits (homologs), GO terms, and InterPro domains assigned to each protein-coding gene have been imported into MongoDB, resulting in more than 126 million records in the database for searching. Indexing of gene/transcript IDs, functional descriptions, GO and Interpro terms, and TF/TR and protein kinase family names has been performed in the database, allowing for efficient search of large amounts of data. The interactive web interfaces have been developed using the Flask web framework and HTML.

## DATABASE FUNCTIONS

### Search interface

To enhance user convenience in searching large-scale genomic data of horticultural crops, we have designed the search page to resemble popular search engines such as Google and Microsoft Bing. Multiple search methods have been streamlined into a single search box, thereby allowing users to search for genes of interest by entering various types of keywords and other related information, without requiring any prior experience or specialized training (Figure 1A). Currently, the keywords could be the name of the species and gene, gene ID, the functional description of the gene, the family name of the transcription factor or protein kinase, or the GO or IPR ID. It is acknowledged that scientific names of crop species may be challenging to enter accurately than common names. Additionally, it is often difficult for users to remember precise information, such as IDs for genes, GO and IPR terms. To address this issue, we have implemented an auto-completion function for entering keywords. This feature prompts users with suggestions based on the information stored in the backend database after entering 2-3 characters, aiding in the accurate entry of information mentioned above. For example, when users search for tomato genetic information, they can use the common name ‘tomato’ or the Latin name ‘*Solanum lycopersicum*’ for the query. When entering the first few characters, the HSE will automatically prompt and complete the corresponding name for users to choose (Figure 1B). After selecting species keywords, users can enter other keywords such as gene ID, gene name, gene functional description, etc (Figure 1C-E).

**Figure 1.**
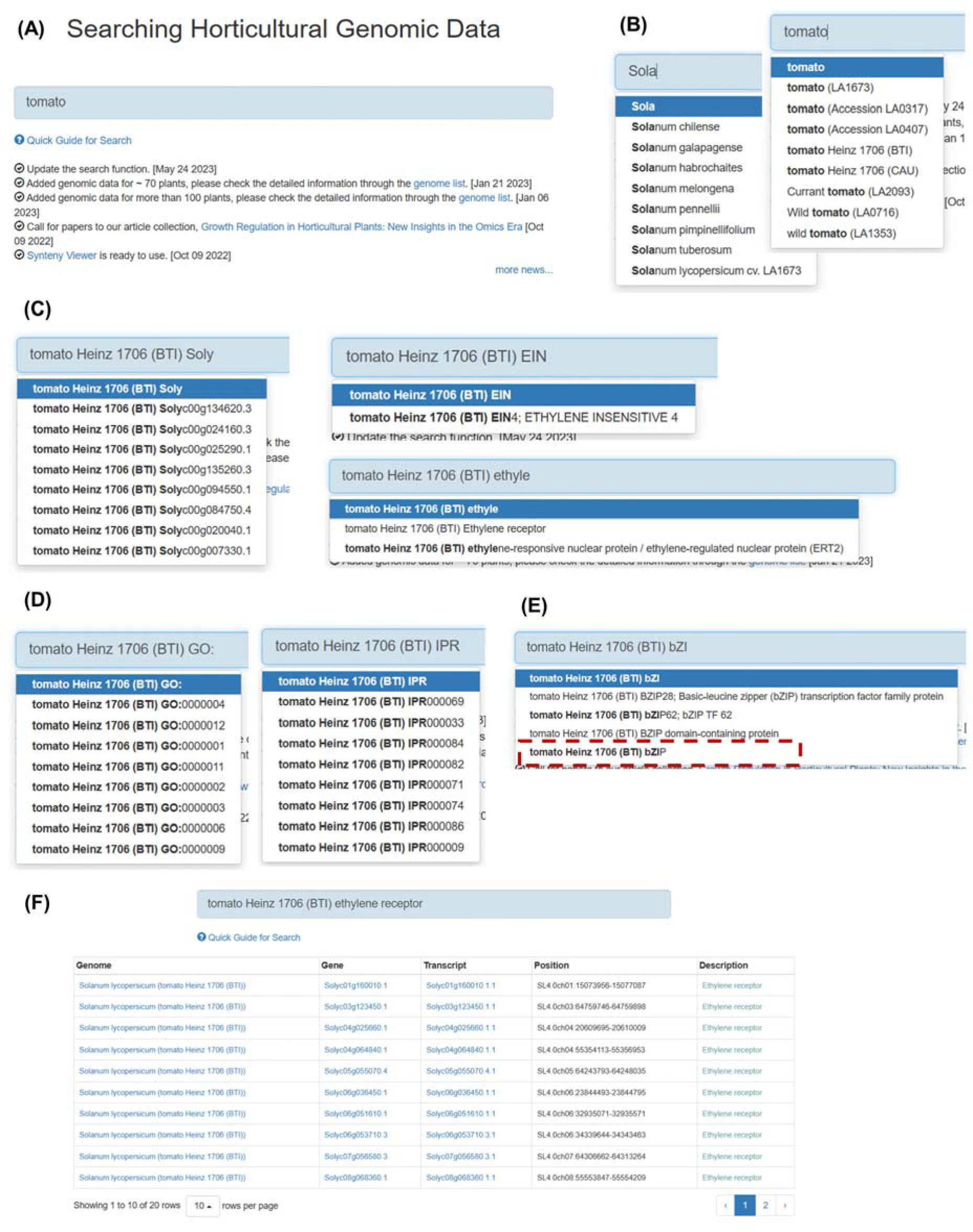
Search interface and result pages in HortGenome Search Engine. **(A-E)** Screenshots of the search interfaces. **(F)** Gene list of search results, including plant name, gene/transcript ID, genomic location and functional description.

The search returns a gene list with the corresponding species name, gene/transcript IDs, gene locations, and gene functional descriptions (Figure 1F). The species name and gene/transcript IDs are linked to the corresponding genome page of the species and gene/transcript pages, respectively. In addition, if user enters a keyword that combines the name of a species and the name of a specific TF/TR/PK family (Figure 1E), the results will directly return to the corresponding gene family page of the species (Figure 3).

**Figure 2.**
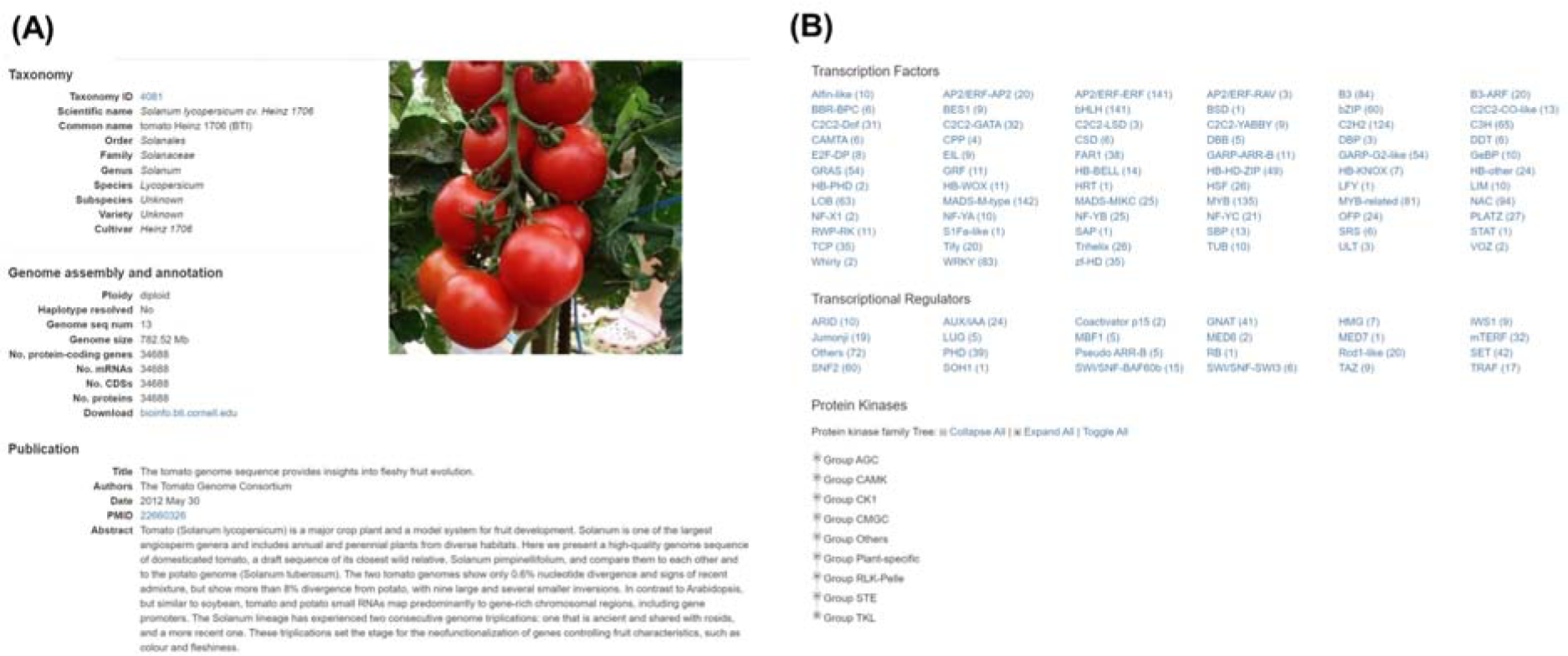
Genome page in HortGenome Search Engine. **(A)** Screenshot of the genome page containing the genome information and picture of the plant. **(B)** Screenshot of the genome page containing transcription factors, transcriptional regulators, and protein kinases identified from the genome.

**Figure 3.**
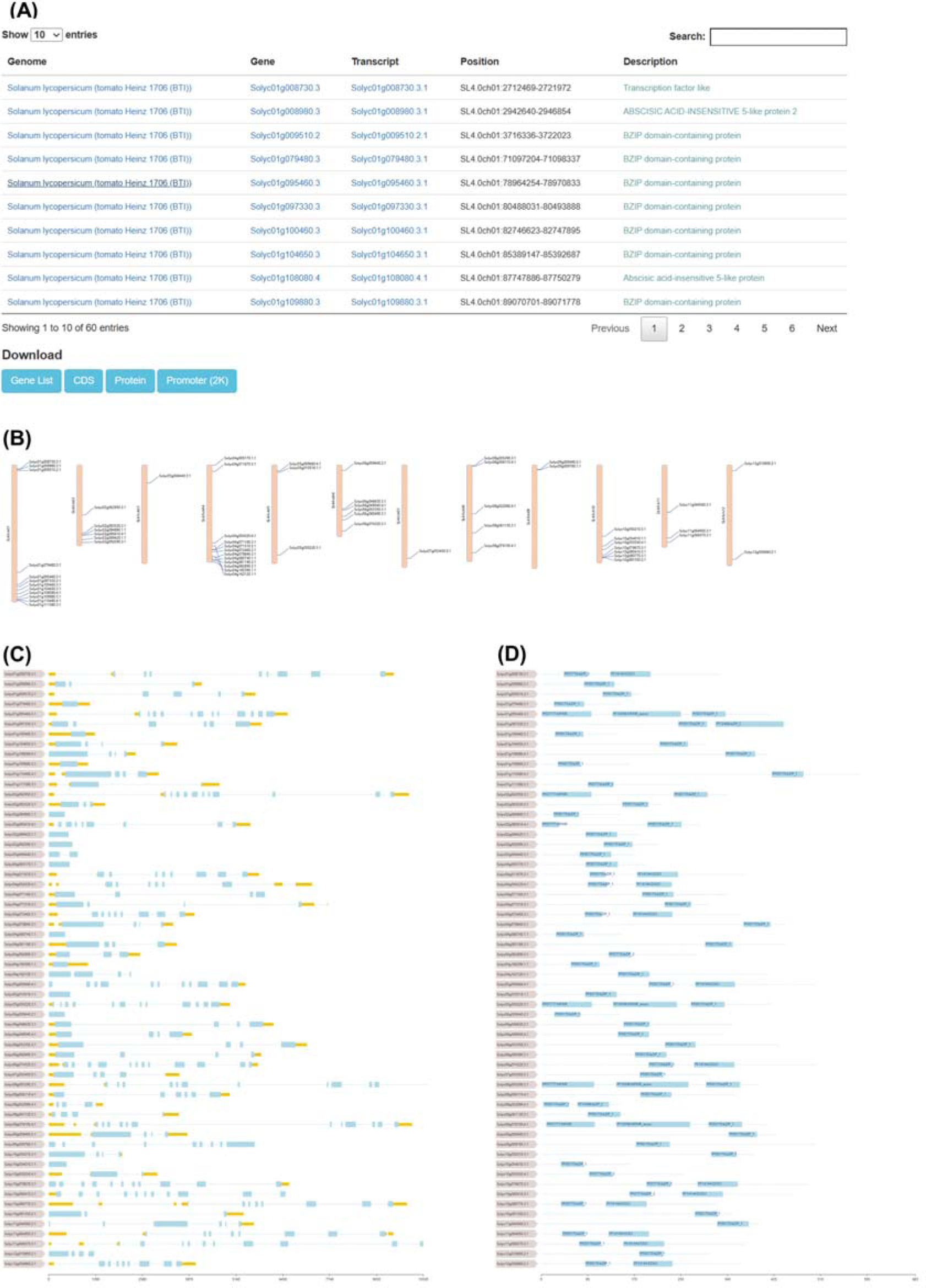
Gene family page in HortGenome Search Engine. Screenshots of the list and download links **(A)**, locations on chromosomes **(B)**, structure **(C)** and functional domains **(D)** of the tomato bZIP family genes.

### Genome page display

The genome page displays basic information about the plant species and the genome assembly, and is comprised of three sections: taxonomy, genome assembly and annotation, and publication. The taxonomy section provides the scientific name, common name, and taxonomy information of the plant species, and the taxonomy ID is linked to the GenBank taxonomy database. The ‘genome assembly and annotation’ section shows the information about genome assembly size, the numbers of genome sequences, genes, mRNAs, CDS, and proteins, as well as the ploidy level information and the download link of the genome assembly. For the publication section, the title, authors, abstract, and publication date of the corresponding genome paper, which were automatically retrieved from PubMed according to the PubMed Identifier (PMID), are displayed (Figure 2A).

On the genome page, an additional pagination is available to display the names and numbers of transcription factors, transcriptional regulators, and protein kinases identified for the selected genome (Figure 2B). Clicking on a family name directs to the corresponding gene family page.

The gene family page displays homologous genes belonging to the same family, as well as gene location, structure, and functional domains. At present, only genes from the transcription factor, transcriptional regulator, and protein kinase families identified by iTAK^22^ can be searched and displayed. For example, searching for the bZIP transcription factor of tomato will display all 60 bZIP genes identified in the genome. The page provides download links to retrieve gene list, CDS, protein, and promoter sequences of these bZIP genes (Figure 3A). The location of genes on chromosomes, gene structure, and protein functional domains are valuable information to study gene families. Therefore, the gene family page of HSE diaplays the images of gene location, structure, and protein functional domains for these homologous genes (Figure 3B-D), which provide convenience for studying the function and evolution of the corresponding gene families.

### Gene and transcript page display

Each gene or transcript has a detailed feature page that contains all the related sequences and annotation information. The gene feature page forms different paginations based on the content types (Figure 4). The overview pagination contains information about plant species, gene ID, location, strand, and functional description, as well as transcripts belonging to this gene. The gene structure is represented by its primary transcript and displayed using FeatureViewer ^26^ (Figure 4A). The sequence pagination contains gene, mRNA (primary transcript), CDS, and protein sequences (Figure 4B). In the BLAST pagination, it shows the top 5 homologs identified from the GenBank, UniProt, and TAIR databases, respectively. The BLAST hit accession IDs are linked to the corresponding databases, which allow users to access the expression, interaction, protein structure, and other information of the homologous genes from other databases. The detailed sequence alignment of the BLAST result is shown in a popup page when clicking the ‘Show’ link (Figure 4C). The domain pagination lists the functional domains identified from the protein sequence of this gene (Figure 4D). The gene ontology pagination lists the GO terms assigned to this gene and the GO IDs are linked to the AmiGO database which provides details of the GO terms (Figure 4E). The TF/TR/PK pagination shows the family name if the gene is identified as belonging to a specific TF/TR/PK family, which is linked to the corresponding gene family page. The syntelog pagination contains the collinear gene pairs and syntenic blocks related to this gene (Figure 4F).

**Figure 4.**
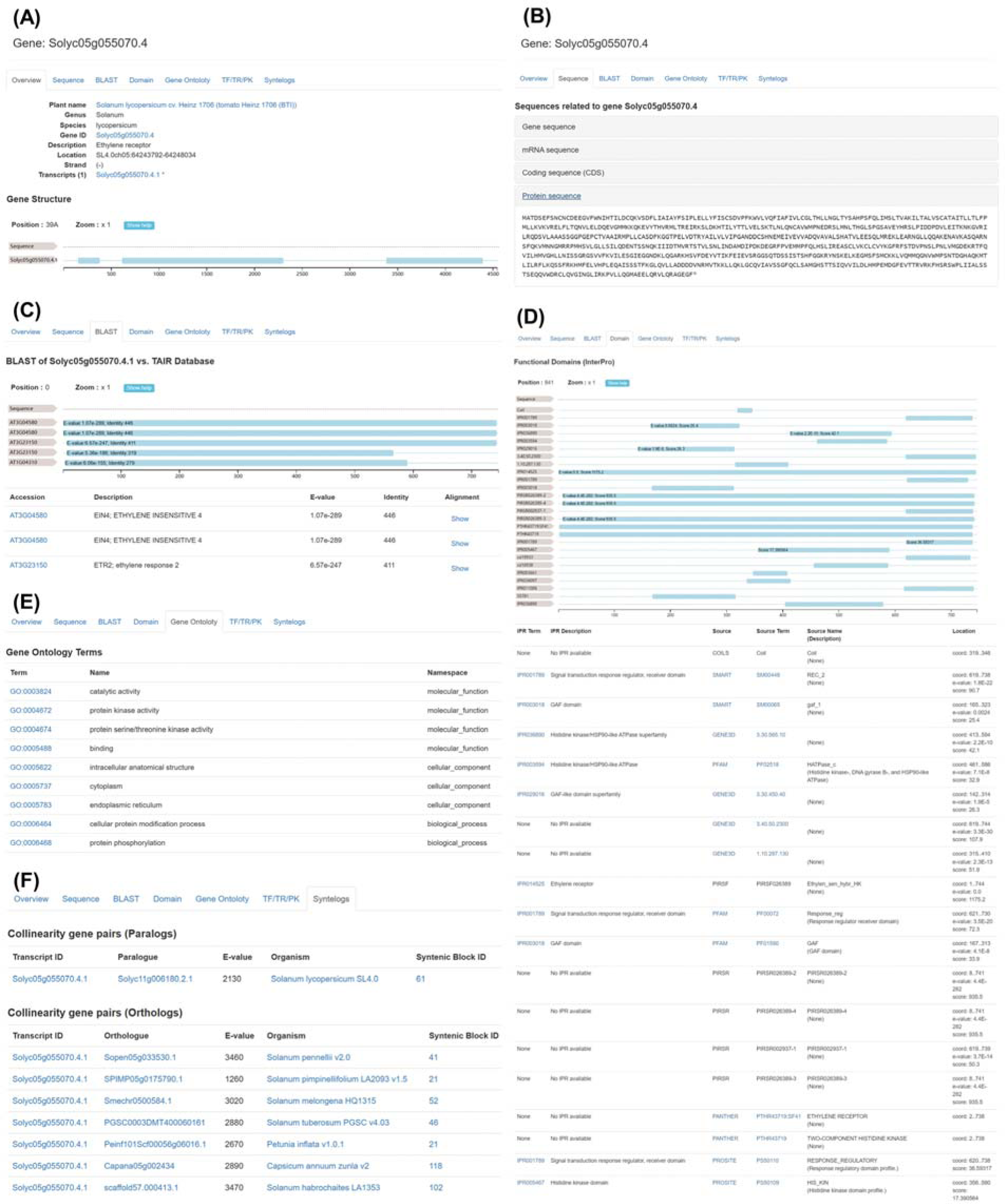
Gene feature page in HortGenome Search Engine. **(A)** Screenshot of the gene page containing basic information and gene structure. **(B)** Screenshot of the gene page containing gene, mRNA, CDS, and protein sequences. **(C)** Screenshot of the homolog genes and sequence alignments from the BLAST results. **(D)** Screenshot of the functional domains predicted from the protein sequence of the gene. **(E)** Screenshot of the GO terms assigned to the gene. (F) Screenshot of the gene page containing collinear gene pairs.

## BLAST

We implemented the online BLAST tool, one of the most widely used tools in genome databases, using the SequenceServer^27^. In the query interface, the indexed genomes are organized in a hierarchical taxonomy display using jsTree (https://www.jstree.com/). The BLAST indexed databases are categorized into nucleotide and protein databases. The nucleotide databases include the BLAST indexes for genome and mRNA/CDS sequences, and protein databases contain all indexes of protein sequences. With this interface, the BLAST search can be performed more flexibly (Figure 5A). For example, by providing a DNA or protein sequence, the user can search against the sequences from a single plant species, or across the entire genus and family, or all plant species in the database. This provides a useful tool for studying gene function and evolution.

**Figure 5.**
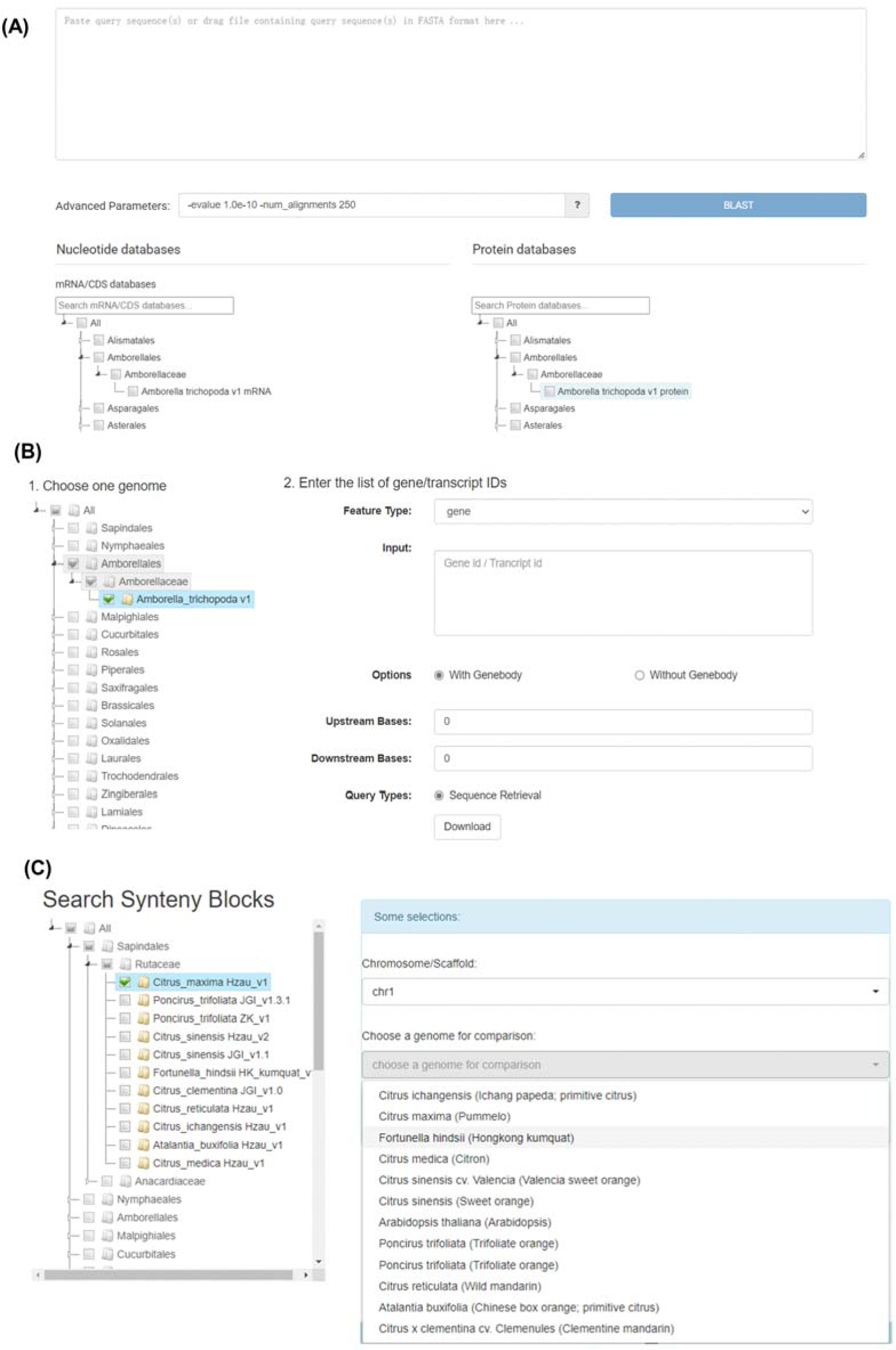
Query interfaces of data mining tools in HortGenome Search Engine. **(A)** Screenshot of the BLAST query page. **(B)** Search interface of ‘Batch Query’. **(C)** Search interface of ‘Synteny Viewer’.

### Batch Query

Genomic and functional genomic studies typically generate large lists of interesting genes, and retrieving nucleotide or protein sequences and functional annotations of these genes for downstream analyses is essential to understand the underlying biological processes. Similar to the online BLAST tool, a hierarchical taxonomy tree is provided in the ‘Batch Query’ interface for easily selecting the genome to be analyzed. The query options will be changed dynamically according to the selected feature type. By selecting the ‘gene’ feature type, sequences containing exons, introns and the upstream and downstream sequences of a list of genes can be extracted (Figure 5B). By selecting ‘mRNA’ or ‘protein’ feature type, in addition to extracting mRNA and protein sequences, the query also allows for retrieving functional descriptions, and family information for TFs, TRs and PKs.

### Enrichment analysis

Genomic and functional genomic analyses are capable of producing extensive lists of genes that are of interest. However, it is crucial to translate these lists into biologically relevant information to gain a deeper understanding of the underlying molecular mechanisms of the related biological processes. Enrichment analysis is a potent method that can be employed to identify classes of genes that are overrepresented in a list of genes. This approach enables the identification of highly dynamical biological processes or biochemical pathways under specific experimental conditions or developmental stages. In order to facilitate the enrichment analysis of gene and transcript data for hundreds of genomes, a hierarchical taxonomy tree has been constructed for the ‘GO Enrichment Analysis’ and ‘KEGG Enrichment Analysis’ tools, utilizing the same structure as that used in BLAST and ‘Batch Query’. The ‘GO Enrichment Analysis’ tool has been implemented through the use of the Perl module GO::TermFinder, which employs the hypergeometric distribution test to determine enriched GO terms^28^. Similarly, the ‘KEGG Enrichment Analysis’ tool has been developed using KEGG pathways assigned to genes via eggnog-mapper, with enrichment significance calculated through the hypergeometric distribution test. The resulting enrichment analysis output page provides a list of enriched GO terms and KEGG pathway names, with links to the relevant GO and KEGG databases^29,30^. Additionally, genes corresponding to each enriched GO term or KEGG pathway are included with links to relevant gene pages in HSE. Overall, GO and KEGG enrichment analyses are essential tools for the interpretation of genomic and functional genomic data, and their use is critical for advancing our understanding of complex biological systems.

### Synteny Viewer

We have previously developed ‘Synteny Viewer’ as an extension module of Tripal to view genome synteny and homologous gene pairs between different cucurbit genomes^9^. The tool has been adopted by many genome databases, including Genome Database for Rosaceae (https://www.rosaceae.org)^8^, ZEAMAP (http://zeamap.com)^31^, etc. In HSE, the ‘Synteny Viewer’ has been re-implemented using Python/FLASK for managing the large amount of comparative genomic data generated from hundreds of plant genomes. To facilitate the search of massive amount of synteny blocks and homologous gene pairs, the genome selection form is designed with genomes well organized through a hierarchical taxonomy tree. The chromosome/scaffold selection drop-down list and the compared genome drop-down list will be automatically updated according to the selected genome (Figure 5C). The search result provides a circos plot that displays synteny blocks for query and compared chromosomes/scaffolds. Each synteny block is linked to a complete list of homologous gene pairs within the block, and each gene is linked to the detailed gene feature page mentioned above.

## CONCLUSIONS AND FUTURE DIRECTIONS

We have developed a universal search engine, HSE, that allows querying genes, functional annotations, and homologous gene pairs for hundreds of genomes of horticultural crops. More than 16 million genes with comprehensive functional annotations as well as 1,832,351 synteny blocks and 413 million homologous gene pairs from 434 genome assemblies are stored in NoSQL document-oriented database for searching. It is worth mentioning that multiple indexes have been established on the document-oriented database to facilitate users to search genes in a more flexible way through a simple search box, which sets HSE apart from other plant genomic databases. Furthermore, several popular data mining tools of genomic databases have been implemented in HSE, including enrichment analysis of GO terms and KEGG pathways, ‘Batch Query’ for retrieving gene sequences and functional annotations, ‘Synteny Viewer’, and BLAST.

We will continue to collect genomic data of horticultural crops for HSE. HSE will be updated every six months or new horticultural genomes are available. In addition, users can submit genomes to HSE by contacting us. In the future, we will expand the scope of data search to cover other omics data such as gene regulatory networks, gene expression, genotype and phenotype. Furthermore, additional online data mining and visualization tools based on the horticultural crop genomes will be implemented in HSE.

## Acknowledgements

We thank Bo Yuan at Lianzhi Technology Co., Ltd. for assistance with computing acceleration.

This work was supported by grants from the Beijing University of Agriculture (Start-up fund) to Y.Z., Young Teachers’ Research and Innovation Capacity Enhancement Program QJKC2022044 and Beijing Municipal Education Commission Scientific Research Plan Project KM202310020010 to S.W. The computing power was supported by the Alibaba Cloud.

## Contributions

Z. Fei and Y. Zheng designed the project. S. Wei, Y. Deng, S. Wu, H. Peng, X. Zhai, S. Zhou, J. Li, H. Li, Y. Feng, Y. Yi, R. Li, H. Zhang, Y. Wang, R. Zhang, L. Ning, and Y. Zheng performed data collection. S. Wei, Y. Qing, H. Peng, and S. Wang performed data analysis. S. Wei and Y. Zheng wrote the code for database construction. Y. Yao, Z. Fei, and Y. Zheng supervised the project and wrote the manuscript. All authors read and approved the final manuscript.

## Data availability statement

All datasets have been made publicly available at http://hort.moilab.net/

## Supplementary data

Supplementary data is available at Horticulture Research online.

